# Structural Basis of σ^54^ Displacement and Promoter Escape in Bacterial Transcription

**DOI:** 10.1101/2023.06.09.544244

**Authors:** Forson Gao, Fuzhou Ye, Bowen Zhang, Nora Cronin, Martin Buck, Xiaodong Zhang

**Author notes:** Lead contact : Xiaodong Zhang.

## Abstract

Gene transcription is a fundamental cellular process carried out by RNA polymerase (RNAP). Transcription initiation is highly regulated and in bacteria, transcription initiation is mediated by σ factors, which recruit RNA polymerase (RNAP) to the promoter DNA region and facilitate open complex formation, where double stranded DNA is opening up into a transcription bubble and template strand DNA is in position for initial RNA synthesis. During initial transcription, DNA downstream of the transcription start site is fed into the active site of RNAP, whilst the upstream promoter DNA remains tethered to the RNAP via the σ factor, resulting in a build-up of tension. In order to progress to the processive elongation state, RNAP must escape from the promoter, and displace or dissociate the σ factor. Bacteria s factors can be broadly separated into two classes with the majority belong to the s70 class, represented by the σ^70^ that regulate housekeeping genes. σ^54^ forms a class on its own and regulate stress response genes. Extensive studies on σ^70^ have revealed the molecular mechanisms of σ^70^ promoter escape while how σ^54^ transitions from initial transcription to elongation is unknown. Here we present a series of cryo electron microscopy structures of the σ^54^ factor with progressively longer RNA, revealing the molecular mechanism of σ^54^ displacement and promoter escape. Our data show that the initial instability is driven by DNA scrunching, and the final displacement steps are driven by both RNA extension and DNA scrunching.

## Introduction

Gene transcription is a fundamental cellular process carried out by RNA polymerase (RNAP). The multi-subunit RNAPs are conserved from bacteria to human, with the active site located at the bottom of the RNAP cleft formed by the two highly conserved large subunits (1) (Fig. 1). Transcription initiation in bacteria involves the recruitment of RNAP by sigma factors (σ) to upstream promoter regions that demarcate the transcription start site (TSS) (2). Bacteria have several σ factors with the majority belonging to the σ^70^ family, represented by the housekeeping σ^70^ (3). In 60% of bacteria, a major variant σ factor, σ^54^, forms a class of its own lacking structural and sequence similarity to σ^70^. In σ^54^ mediated transcription initiation, σ^54^ recruits RNAP to specific gene promoters responsible for regulating a variety of stress responses, including nutrient depletion, membrane stress, antibiotic exposure and biofilm formation (4). Transcription initiation involves several distinct steps including promoter search and subsequent closed complex formation, when RNAP-σ holoenzyme stably engages with promoter DNA, which remains duplexed and outside of RNAP active site. The closed complex is subsequently converted to an open complex, when promoter DNA is opened up into a transcription bubble and the template strand is delivered into RNAP active cleft, ready for transcription (5).

**Fig. 1.**
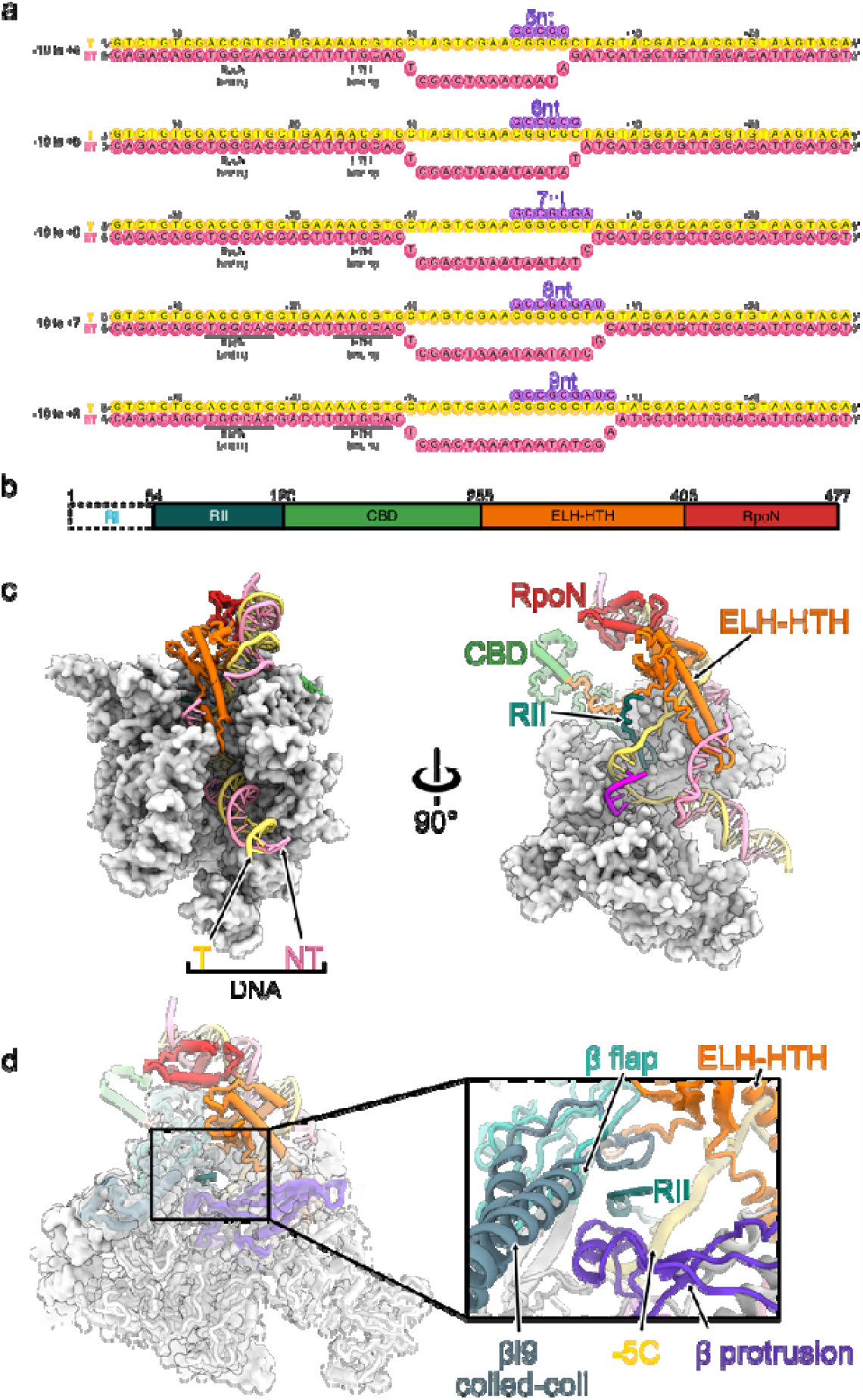
Overall structure and nucleotides used in this study. a) Design of the DNA-RNA scaffolds used in this investigation, showing complete DNA nucleotide sequence. b) Overview of the σ^54^ domain structure. RI is shown in white, as it is unresolved. c) Left: overview of 5 nt initial transcribing complex (RP5nt) structure, viewed from the downstream. RNAP is shown in white. Right: cross-section of RP5nt, with RII highlighted, and ⍰ and β subunits hidden. d) RII entrance into the RNAP core. Right panel: Overview of the β face of the 5nt pre-translocated elongation complex. Left panel shows a zoom in showing the RII exiting from the <u>β</u> protrusion, <u>β</u> flap, <u>β</u>i9 coiled-coil and HTH subdomains.

Previous studies, including single molecule biophysical studies and structural studies, established that during initial transcription, upstream DNA remains bound by RNAP-σ while DNA downstream of TSS is brought into RNAP channel, thus resulting in an enlarged transcription bubble inside the RNAP catalytic site (6-8).

Promoter escape is the final step in transcription initiation when RNAP is released from the promoter region and translocates downstream while the enlarged transcription bubble collapses down to the ∼10 nt bubble in the elongation complex. During elongation, the transcription bubble is maintained by a series of conserved loops in the RNAP core (Supplementary Fig. 1, discussed in more details in (9)). The DNA-RNA hybrid is cordoned between the β’ bridge helix and the β’ lid. At the β’ lid and β fork loop 1, the 5’ end of the RNA is diverted into the RNA exit channel, made up of the β’ zipper, β flap and β switch 3 loops. The upstream and downstream edges of the transcription bubble is maintained by the β’ rudder and β fork loop 2 respectively.

σ displacement is essential in enabling bacterial RNAP to be released from the promoter and for processive RNA synthesis during elongation (10).

σ^70^ and σ^54^ are structurally and mechanistically distinct, although contain subdomains serving analogous functions (Fig. 1b) (11). Both σ factors contain flexible linkers that descend into the catalytic site of the RNAP and occupy the RNA exit channel (Fig. 1c, supplementary Fig. 1). In the RNAP-σ^70^ holoenzyme, region 1.1 occupies the downstream DNA site, whilst region 3.2, also known as the σ finger, occupies the RNA exit channel, with region 4.1 sitting on the RNA exit site(12, 13). In the RNAP-σ^54^ holoenzyme, region II (RII) occupies both the downstream DNA binding site and the RNA exit channel, with the core binding domain (CBD) sitting on the RNA exit site (11, 14). The CBD interacts with the RNAP ⍰-CTD, β flap, β’ zinc finger (11).

Previous studies have revealed the mechanism of σ displacement in σ^70^, as either arising from RNA extension causing the 5’ end of the RNA to push against the σ finger (15, 16), or DNA scrunching as a result of an enlarged transcription bubble (6, 7, 17). Given the differences between σ^54^ and σ^70^, it is unclear how σ^54^ displacement occurs during initial transcription that leads to elongation. In this study, we used single particle cryo electron microscopy (cryoEM) to determine the structures of initial transcribing complexes, which allow us to propose the mechanism of σ^54^ RNAP promoter escape.

## Results

To capture structures of initial transcribing complexes that lead to promoter escape, we designed DNA-RNA scaffolds based on the well characterised nifH promoters with increasing sizes of transcription bubble and RNA lengths to mimic the growing mRNA length and the corresponding enlarged transcription bubble (Fig. 1a)(14). Using cryoEM and single particle analysis, (Figs. 1-2, Supplementary Figs. 2-5, Tables 1-2). We determined six structures of RNA- and σ^54^-bound initial transcription complexes containing 5 nt to 9 nt RNA.

**Fig. 2.**
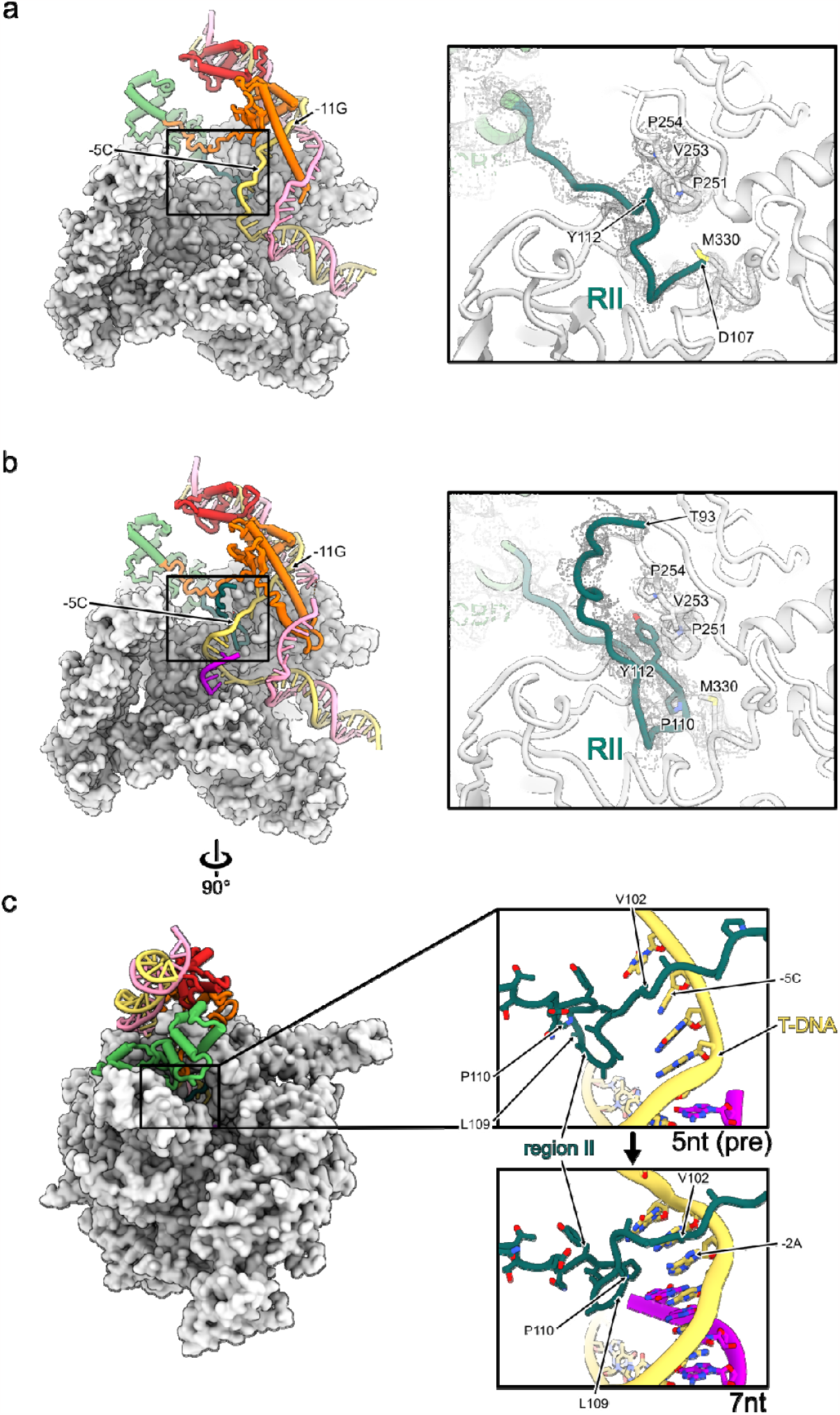
Region II tip and transcription bubbles in RPitc. (a) RII tip is only partially resolved in the open complex (RPo) (b) in the 5 nt initial transcribing complex (RP5nt, pre-translocated), RII tip is resolved and kinks sharply at the site close to the template strand DNA. (c) Template strand DNA interacts with RII tip. Left panels Cross section of the complexes Right panels zoomed in regions highlighted by black box

**Table 1:** Data collection parameters

| Sample | Pixel Size<br>(Å/pixel) | Total Dose<br>(e<sup>-</sup>/Å<sup>2</sup>) | Defocus Range<br>(μm) | Total Movies |
| --- | --- | --- | --- | --- |
| RP5nt | 1.1 | 30 | -3 to -1 | 12 309 |
| RP6nt | 1.06 | 40 | -2.7 to -1 | 9188 |
| RP7nt | 1.06 | 30 | -3 to -1 | 4185 |
| RP8nt | 1.072 | 40 | -2.7 to -1 | 11 198 |
| RP9nt | 1.06 | 30 | -3 to -1 | 6121 |

**Table 2.** Model validation statistics for longer transcript complexes

| Sample | RP5nt pre-translocated | RP5nt post-translocated | RP6nt | RP7nt | RP8nt | RP9nt |
| --- | --- | --- | --- | --- | --- | --- |
| Refinement |  |  |  |  |  |  |
| Model composition |  |  |  |  |  |  |
| Non-hydrogen atoms | 29191 | 28821 | 28681 | 28965 | 28878 | 28605 |
| Protein residues | 3668 | 3668 | 3638 | 3635 | 3617 | 3617 |
| Nucleic acid residues | 102 | 100 | 101 | 104 | 105 | 103 |
| Ligands | 2 Zn<sup>2+</sup> | 2 Zn<sup>2+</sup> | 2 Zn<sup>2+</sup> | 2 Zn<sup>2+</sup> | 2 Zn<sup>2+</sup> | 2 Zn<sup>2+</sup> |
|  | 1 Mg<sup>2+</sup> | 1 Mg<sup>2+</sup> | 1 Mg<sup>2+</sup> | 1 Mg<sup>2+</sup> | 1 Mg<sup>2+</sup> | 1 Mg<sup>2+</sup> |
| B factors (Å<sup>2</sup>) |  |  |  |  |  |  |
| Protein | 63.87 | 40.32 | 63.05 | 50.02 | 42 | 37.68 |
| Nucleic acid | 23.49 | 95.31 | 101.33 | 101.62 | 156.55 | 180.21 |
| Ligands | 69.42 | 27.99 | 58.31 | 54.39 | 33.57 | 46.53 |
| R.m.s. deviations |  |  |  |  |  |  |
| Bond lengths (Å) | 0.004 | 0.003 | 0.005 | 0.005 | 0.005 | 0.003 |
| Bond angles (°) | 0.679 | 0.604 | 0.71 | 0.765 | 0.681 | 0.643 |
| Validation |  |  |  |  |  |  |
| MolProbity score | 1.94 | 1.85 | 1.99 | 1.96 | 1.95 | 1.95 |
| Clashscore | 10.87 | 8.71 | 10.84 | 11.06 | 10.12 | 11.03 |
| Poor rotamers (%) | 0.59 | 0.27 | 0.42 | 0.37 | 0.37 | 0.34 |
| Ramachandran plot |  |  |  |  |  |  |
| Favored (%) | 94.29 | 94.37 | 93.3 | 94.13 | 93.59 | 94.28 |
| Disallowed (%) | 0 | 0.03 | 0.03 | 0.03 | 0.03 | 0.08 |

### Structures of RPitc with 5nt mRNA reveal the passage of region II through the RNAP core and interactions with the transcription bubble

Using a combination of focused classification and electron density subtraction, we obtained several distinct structural states within the same dataset, including pre-, post-translocated RNA and σ^54^-bound states, as well as σ-released states (Supplementary Fig. 4). The pre- and post-translocated states were resolved to resolutions of 2.8 and 3.4 ⍰ respectively based on gold standard FSC curves (Supplementary Fig. 3) (18).

Compared to the open promoter complex RPo, the RNAP structure remains largely unchanged (Fig. 2)(14). σ^54^ domains also remain in similar positions except for σ^54^ RII. σ^54^ RII connects region I (RI) and the CBD of the region III (RIII) that sits and blocks the RNA exit channel (Fig. 1b-c) (11). As with the previous RPo and RPitc (4 nt) structures (14), we could only resolve parts of RII (Fig. 2a-b). Residues C-terminus of Y112 are in similar conformations in the RPitc compared to RPo but the tip of RII (residues 107-112) alters the conformation in RPitc compared to RPo (Fig. 2).

For RPo, we could resolve from D107 onwards (Fig. 2a). In the RPitc 5nt structures, we could resolve more residues, to T93, enabling us to trace the path of the RII C⍰ backbone, which traces the RNA exit channel and leads towards the RNA cleft (Fig. 2b). Interestingly this is different from the traces of RII in the RNAP-σ^54^ holoenzyme, where this part of RII occupies the template strand position, just above the bridge helix (Supplementary Fig. 6a)(11). Presumably upon open complex formation, RII relocates to make space for the transcription bubble. In RPitc 5nt, this part of RII now occupies the RNA exit channel (Supplementary Fig. 6a). The N-terminal half of RII is constrained in a narrow tunnel formed by the β-flap, βi9 coiled coil, β-protrusion and an antiparallel β-sheet on extra-long-helix helix-turn-helix (ELH-HTH) (Fig. 1d). This is indeed similar to the RNA exit tunnel (Supplementary Fig. 1). RII contacts the template strand at the -5 position (Fig. 2c), and threads around the template DNA-RNA hybrid and forms a sharp kink at P110, before exiting via the RNA exit channel and connecting to the CBD (Fig. 2b). Indeed, the bases of the template strand interact with RII (Fig. 2c). We now refer to this kink (residues 104-113) as the RII-finger due to its functional similarity with the σ^70^ finger, also known as R3.2 in σ^70^. The RII kink is stabilised by hydrophobic interactions between P110 and Y112 of RII, and the conserved P251, V253 and P254 on the β’ lid and M330 on β’ switch 2 region (Fig. 2b).

### RNA extension causes folding back of the RII-finger

Data collected on each of the initial transcribing complexes (Fig. 1a) were processed using similar strategies as for the RPitc-5nt dataset. In all the datasets, we found a subset of σ^54^-free states, as well as RNA-bound, post-translocated, σ^54^-bound states (Fig. 3a, Supplementary figures 4-5).

There is little difference in RII between 6 nt and 5 nt and indeed RNA remains short of reaching RII (Fig. 3a-b). However, at 7nt RNA, the RII-finger adapts a different conformation in order to accommodate for the growing DNA-RNA hybrid and prevent steric clashes; the tip of the RII-finger is coiled backwards by 9.5 ⍰ (as measured from the C⍰ atoms with the tip of the RII finger). RII tip contains hydrophobic residues L109, P110 and V111 which form hydrophobic interactions with the 5’ base of the RNA (Fig. 2c). In 8nt and 9nt complexes, the density for the RII-finger is partially lost (Fig. 3b), indicating this region becoming flexible and dynamic in nature and the interactions observed in 7nt are lost. In addition, P110 and Y112, observed in 5nt – 7nt complexes, were also not resolved, indicating that these interactions were no longer present. These results suggest that the folded back conformation observed in 7 nt complex is stabilised by specific hydrophobic interactions, therefore likely represent a kinetic barrier during initial transcription.

**Fig 3.**
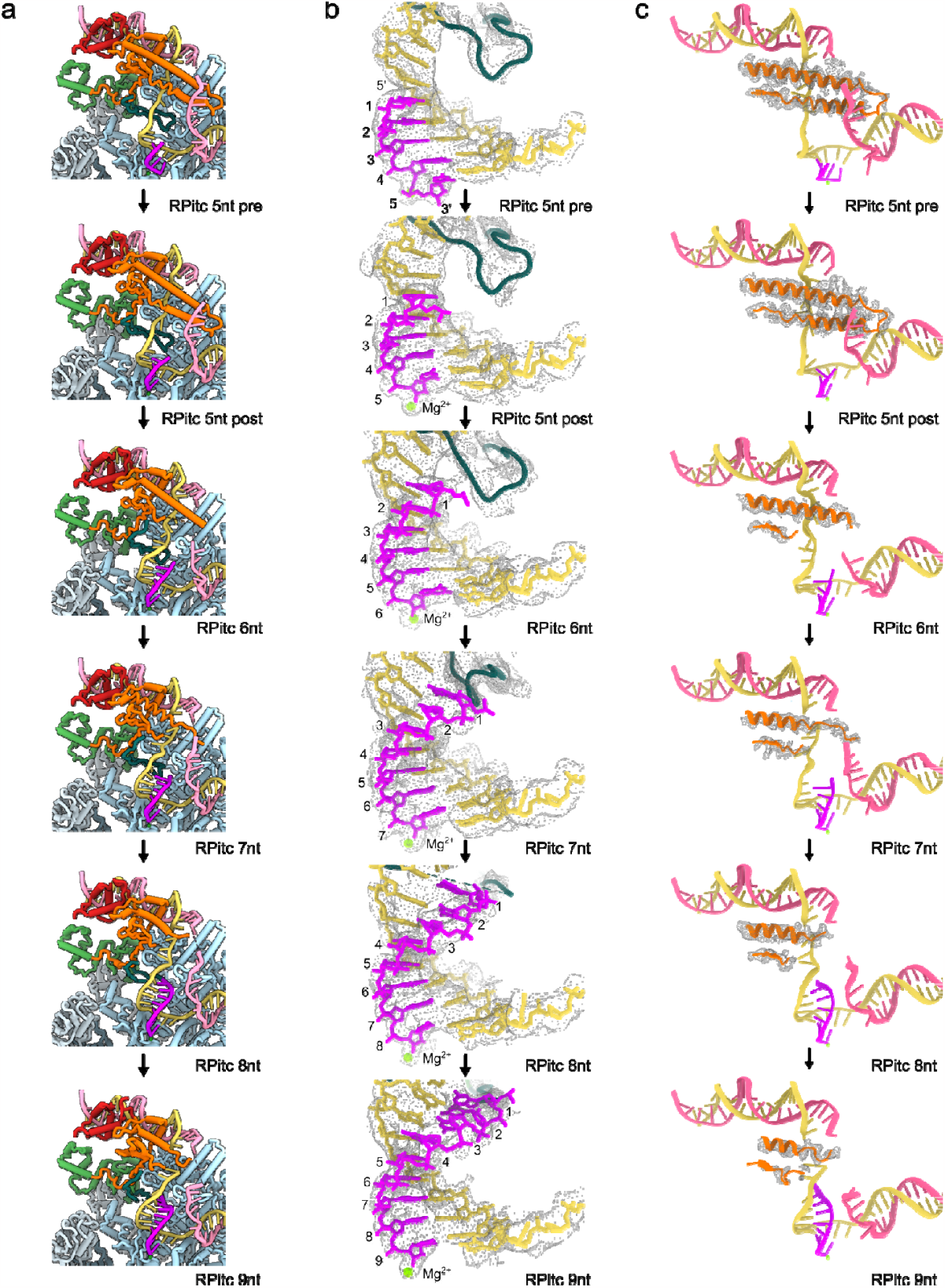
Structural changes during initial transcription: a) overall structures of the initial transcribing complexes from 5 nt to 9 nt with β’ subunit hidden. (b) RNA, template DNA and RII densities and conformational changes at different transcript lengths. At 7nt, RII begins to fold back to accommodate the enlarged DNA-RNA hybrid. (c) Increased RNA leads to loss of ELH density.

Taken together, data presented here suggest that RNA extension to 7 nt causes the folding back of the RII-finger which is stabilised by specific interactions between RNA, DNA and RII-finger. Further RNA extension pushes the RII-finger towards the RNA exit site by breaking the hydrophobic interactions between the β’ lid and RII residues P110 and Y112.

### RNA extension drives DNA scrunching, altering interactions between templates strand DNA and RII-finger

In RPo, the transcription bubble is separated by ELH and the transcription bubble is stabilised by interactions with β and β’ subunits (Fig. 2a) (14). During initial transcription, as in RPitc-5nt, DNA scrunches with the template strand expanding into the back of the cleft and non-template strand, just upstream from the active site nucleotide scrunching towards the β subunit (Supplementary Fig. 1d). The template strand now interacts with RII-finger and with β’ lid and β’ zipper (Fig. 2c, Supplementary Fig. 1). In RPitc-6nt and RPitc-7nt, the template strand (−5 and -6) is scrunched further around RII-finger, β’ lid and zipper, and σ^54^ ELH (Fig. 3, supplementary Fig. 1d) while the non-template strand scrunch around the ELH.

At RPitc-8nt and 9nt, both template and non-template strands scrunch around the ELH, resulting in reduced density for both DNA and the ELH (Fig. 3c). Our results thus show that DNA scrunch is a major mechanism during initial transcription up to 9 nt with progressively more scrunches occurring initially close to the active site and then move upstream towards the ELH. The interactions with β’ lid and zipper play key roles in stabilising the template strand.

### RNA extension increases the dynamics of transcription bubble and σ^54^ ELH

In addition to RII-finger folding back and DNA scrunch, we also observe reduced density for ELH and the transcription bubble surrounding it as the transcription bubble and DNA-RNA hybrid size increase. The ELH density is more smeared from 6 nt of RNA compared to that of 5 nt RNA, indicating that there is an increased flexibility of ELH from 6nt (Fig. 3c). This coincides with the less well-defined density for the non-template strand around here. Indeed, during initial transcription, apart from minor scrunching at the -1 position (nucleotide immediate upstream from the synthesizing site), the non-template strand scrunching mainly occurs around ELH (Fig. 3).

In RPo, ELH separates the transcription bubble between -11 and -7 (Fig. 2a). Apart from the interactions with DNA, ELH has few interactions with the rest of RNAP in the cleft, suggesting that ELH and DNA transcription bubble stabilise each other via their direct interactions. ELH flexibility thus can in part be a result of the enlarged transcription bubble, and the subsequent reduced interactions and constraints between the transcription bubble and ELH. The less constrained ELH will be able to slide and potentially retract out from the transcription bubble. Furthermore, we also observe the loss of resolution of the upstream DNA and upstream DNA binding domains on σ^54^ (RpoN, ELH-HTH) (Supplementary Fig. 2), suggesting that the σ^54^ upstream DNA binding subdomains and DNA are less stably engaged with RNAP with increased RNA lengths.

Taken together, the results presented here suggest a potential path for σ^54^ dissociation. DNA scrunching results in reduced interactions with ELH-HTH, thus allows the upstream DNA-binding RpoN become free from RNAP, and the ELH being released from the enlarged DNA bubble. RNA extension drives RII out of the RNA exit channel, likely pulling RI along with it, and finally dissociation of the CBD from the RNAP core (Fig. 4).

**Figure 4:**
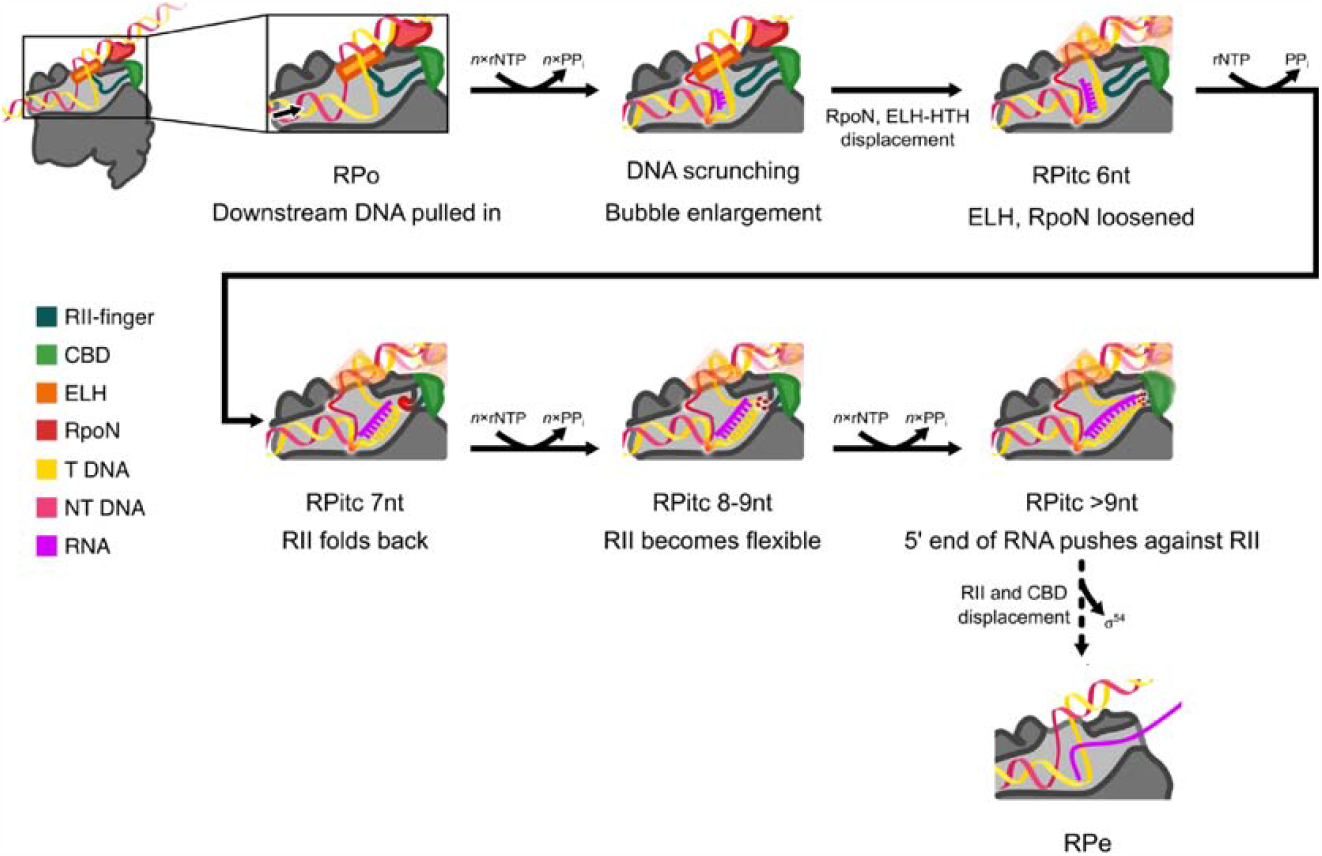
Proposed mechanism of σ^54^ displacement. In RPo, transcription bubble is separated and stabilised by ELH as well as conserved regions in RNAP. Initial transcription results in local DNA scrunching while the structure remains largely unchanged. From 6 nt, DNA scrunching reduces interactions with ELH, enabling ELH and DNA binding domain to be more flexible. At 7 nt, RII-finger folds back and is stabilised by specific interactions with template strand DNA, RNA and RNAP. From 8 nt, RII-finger is no longer stabilised, enabling growing RNA to extend in the exit channel, eventually leading to displacement of RII. ELH and DNA binding domains become more flexible, eventually leading to promoter escape.

## Discussion

### σ^54^ promoter escape results from DNA scrunching and RII conformational changes

The structures of RPitc complexes here show that during initial transcription, the downstream DNA is pushed into the catalytic site, the transcription bubble enlarges. Up to 6 nt RNA, the upstream DNA remains stably bound to the RNAP-σ holoenzyme. This confirms models based on previous FRET experiments using σ^70^, demonstrating RNAP remains bound to the upstream fork of the transcription bubble whilst downstream DNA is pulled in (6). From 7 nt, ELH and upstream DNA becomes more flexible, indicating a global conformational change occurring with the transcription complex.

The ELH interacts with and stabilises the transcription bubble in the RPo structure (14). During initial transcription (from 6 nt), the increased transcription bubble releases the interactions between the bubble and ELH. This increases the flexibility of the ELH, enables it to move out of the bubble. The movement of ELH, and DNA supercoiling caused by scrunching could lead to reduced interactions between the HTH and RpoN subdomains of σ^54^ and the RNAP, as demonstrated by the reduced resolution of these subdomains and increasing amount of elongation complexes observed in the datasets with increasing mRNA lengths.

Given the extensive interactions between σ^54^ and DNA and the reduced interactions with RNAP in the complex with longer RNA, it is possible that the attachment of σ^54^ to upstream DNA (−24 and -12) helps RNAP to be released from σ^54^ while translocating downstream, releasing the tensions of scrunched DNA, transitioning into the elongation complex. However, the conformational changes in σ^54^ upon RNAP dissociation, in particularly around RII and ELH, could also reduce σ^54^ attachment to promoter DNA.

σ^54^ CBD does not interact with DNA directly, and we did not observe significant movement of the β’ rudder, β’ lid, β’ coiled-coil, β’ zipper, β fork loop or β flap during initial transcription. This suggests that transcription bubble enlargement and DNA scrunching is unlikely to significantly perturb the structure of the CBD. However, RII movement, movement of other σ^54^ domains, and elongating RNA or RNAP translocating downstream will ultimately lead to CBD displacement from RNAP, helping with σ^54^ dissociation from RNAP. The linker between σ^54^ RI and RII during initial transcription remains unresolved; the N-terminus of RII enters the catalytic site from the RNAP β side, in between the β protrusion and σ^54^ HTH.

RI is the major inhibitory domain that prevents open complex formation. RI is not resolved in these initial transcribing complex structures. From our previously determined closed and intermediate complex structures, one RI helix is shown to be stabilised by interactions with the 12/-11 promoter DNA region as well as hydrophobic and salt bridge interactions with the ELH, thus constraining ELH conformation and together form an obstacle for DNA loading (19, 20). Recently, RI N-terminal peptide is shown to be remodelled by the bEBP via its ATPase activity and the RI helix is proposed to be unfolded upon ATP hydrolysis of bEBP, releasing its constraints on ELH (20). In these initial transcribing complexes, RI has indeed already been removed from ELH and is now flexible and does not impose any additional constrains on σ^54^ displacement.

Interestingly, despite the important roles we have identified in promoter escape, RII is the least conserved regions in σ^54^ both in sequence identity and sequence lengths, with some species having a significantly shortened RII (for example in Rhodobacter capsulatus) (21). It is therefore possible that in these species, σ^54^ displacement occur at longer RNA lengths, and involve more DNA scrunching, similar to those observed in σ^H^ (ECF) factor (15). Furthermore, the specific interactions with template strand would suggest the dynamics and kinetics of σ^54^ displacement could be promoter-dependent. Indeed, FRET experiments in σ^70^ (Wang et al. XXX) showed that promoter-escape and kinetics are promoter-dependent.

### Comparisons with σ^70^ promoter escape

In σ^70^, as observed in X-ray crystallographic structures, the σ finger is seen to start to fold back in a stepwise fashion from 5 nt RNA (22). In σ^54^, we observe RII-finger folding back at 7 nt RNA to accommodate the increasing DNA-RNA hybrid. As RNA length increases to 9 nt, RII becomes more flexible and disordered. Our studies thus demonstrate that σ-finger refolding is likely a common mechanism for σ displacement, with R3.2 of σ^70^ being functionally similar to σ^54^ RII, and the CBD of σ^54^ playing similar roles to R4.1 of σ^70^ (15, 23). As the RNA extends, the RII-finger being pushed out, CBD dissociates and the rest of RII would be pulled out of the active cleft (Fig. 4). Furthermore, scrunching of the DNA has been previously observed in FRET experiments with σ^70^ and has been shown to also play a role in the build-up of stress within the walls of the RNAP catalytic site. We propose that scrunching of DNA in σ^54^ also plays a significant role, primarily in the release of RII-finger, ELH-HTH and RpoN from the upstream DNA. Using labels directly on the σ-finger, single molecule FRET studies suggest promoter-dependent sigma displacement in σ^70^ (Wang et al., XXXX). The interactions of RII-finger and template strand would suggest σ^54^ displacement is also promoter sequence -dependent.

Despite the functional and mechanistic similarity, there are differences between σ^54^ and σ^70^. R3.2 of σ^70^ enters deeper into the catalytic site compared to RII of σ^54^, consequently it starts to fold back at 5 nt RNA while this only occurs at 7 nt for RII of σ^54^ (Supplementary Fig. 6b). Furthermore, DNA scrunching and RNA extension reduce interactions between DNA and RNAP and σ^54^ and RNAP, while the main interactions between σ^54^ and DNA remain unaffected. It is thus possible that promoter escape involves the translocating RNAP downstream while σ^54^ remains tethered to upstream -24 and -12 regions. On the other hand, σ^70^ core-binding domain R2 is not involved in DNA interactions or RNA extension, it is therefore consistent with σ^70^ sometimes remaining RNAP bound after promoter escape (24, 25).

## Acknowledgements and funding sources

We are grateful to Paul Simpson from the centre for structural biology (CSB) at Imperial College for help and support during initial sample screening and sample optimisation. Data were collected at eBIC, Diamond Light Source and LonCEM. This project is funded by the UKRI to X.Z. and M.B. (BB/M011178/1). F.G. is funded by a BBSRC DTP studentship.

## Materials and Methods

### Protein Purification

Protein purification was carried out as previously described, using a R336A bypass mutant for open complex formation in the absence of activator protein (14). RNAP-σ^54^ _R336A_ was formed by incubating RNAP with σ^54^ _R336A_ in a 1:4 molar ratio at 4 ⍰ for 1 hour, before gel filtration using a Superose 6 10/300 column (GE Healthcare) equilibrated with GF buffer (20 mM Tris-HCl pH 8, 150 mM NaCl, 5% v/v Glycerol, 2 mM TCEP).

### Design of DNA-RNA scaffolds

DNA and RNA were synthesized as single stranded oligos by IDT and the oligos were annealed by mixing equimolar amounts of complementary strands in 20 mM Tris, pH 8.0 and heating to 95 °C for 2 minutes before cooling to 4 °C, by reducing 2 °C per minute. Oligos were used directly in cryoEM sample preparations.

### Sample preparation

17 μM RNAP-σ^54^ _R336A_ was incubated with 18.7 μM of DNA-RNA scaffold in the presence of buffer EM1 (20 mM Tris-HCl, 150 mM NaCl, 10 mM MgCl2, 1 mM TCEP) for 1 hour at 4 °C. Following incubation, samples were buffer exchanged into Buffer EM2 (20 mM Tris-HCl, 150 mM KCl, 5 mM MgCl2, 5 mM TCEP) using a 0.5 ml Zeba™ 7K MWCO desalting column as per the manufacturers’ protocol. 8 mM CHAPSO was then added immediately before cryoEM grid preparation.

### Grid preparation

300 mesh holey gold C-flat R1.2/1.3 grids (ProtoChips), were plasma cleaned in air for 30 seconds (Harrick Plasma). 4 μl of complex was deposited onto plasma cleaned grids. The blotting parameters were as follows: wait time 30 seconds, blot time 2 seconds and blot force -8. Grids were made using a Vitrobot™ Mark IV (FEI) at 4 °C and 100 % humidity with Grade 595 Vitrobot™ filter paper (Electron Microscopy Sciences). All grids were plunge frozen using liquid ethane and stored in liquid nitrogen.

### Data Collection

Datasets were collected on a Titan Krios (ThermoFisher Scientific), operated at 300 kV, with a K3 direct electron detector and a Bioquantam energy filter (Gatan). Movies were collected at a nominal magnification of 81,000× and a slit width of 20 eV (Error! Reference source not found.). Data collection was carried out using EPU software (FEI).

### Image processing

All image processing was carried out in RELION 4.0 (26), using the built-in MOTIONCORR implementation in RELION (27) and CTFFIND4 (28) with particles picked using Topaz (29). For RPitc 5nt complexes, the published RPitc complex was used as an initial reference (EMDB: 4397) (14), whereas the other datasets used the 1 nt shorter complex as a reference. Classes were selected based on definition of key σ^54^ subdomains using a combination of focused 3D classification and subtraction and recentring. A range of masks were tested, with the most accurate angular assignments coming from masks of CBD and RII to find σ^54^-bound classes within the dataset, with tighter masks around the DNA-RNA hybrid used in later stages to identify RNA bound complexes.

### Model building and Refinement

All structural models were built using COOT (30) and refined using real-space refine in PHENIX (31, 32). All figures were prepared using UCSF ChimeraX (33).

### Quantification and Statistical Analysis

Please see Table 2 for quality of 3D reconstructions and models.

