## Supplementary figures for "Structural Basis of σ^54^ Displacement and Promoter Escape in Bacterial Transcription"

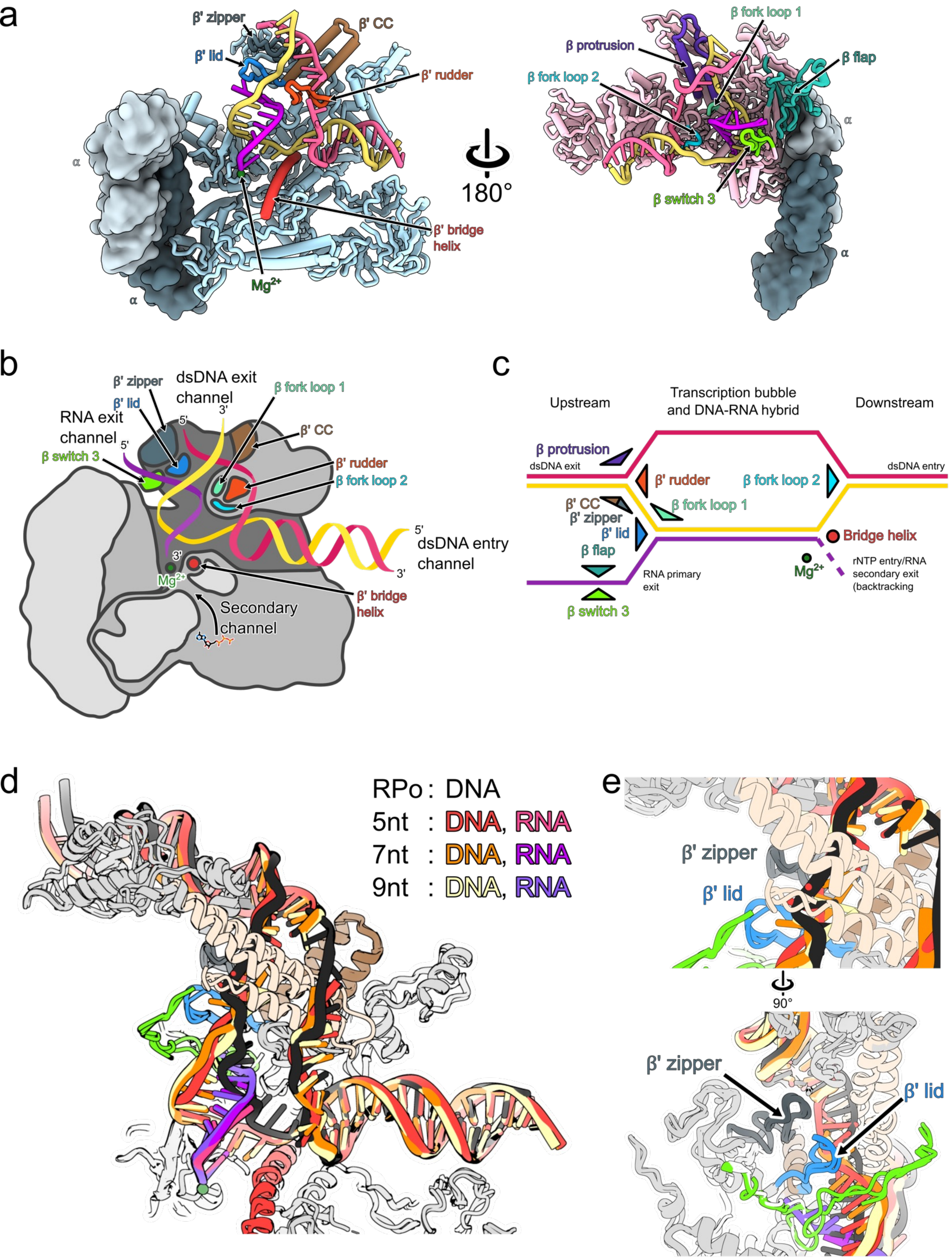


Supplementary Figure 1: Key conserved features in RNAP. a-c, elongation structure showing regions responsible for maintaining the transcription bubble. Adapted from PDB 6XDR. d, maintenance of the transcription bubble in the initial transcribing complex.

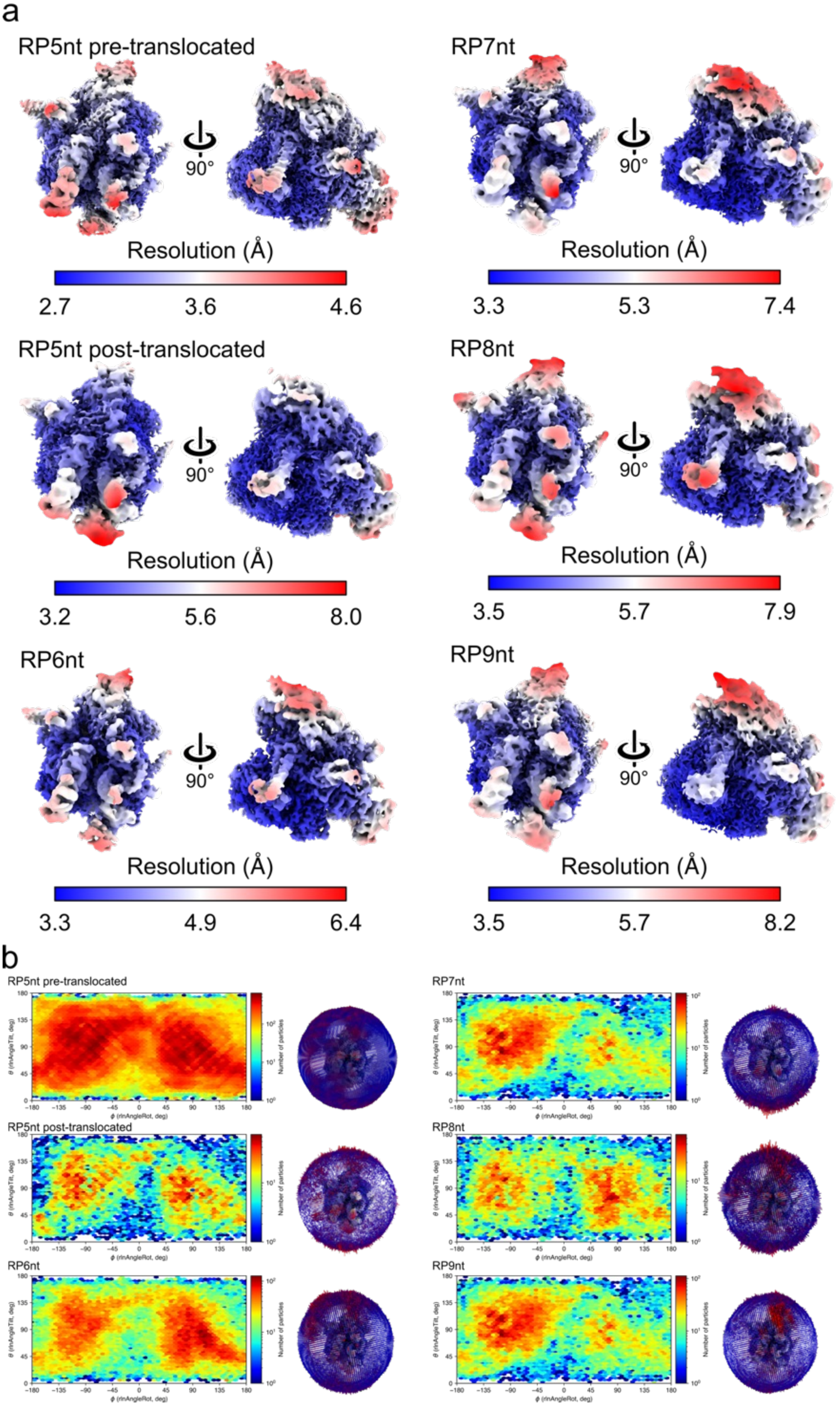


Supplementary Figure 2. Resolution maps, angular distribution of contributing particles: a) Local resolution maps and b) Angular distribution maps of the final reconstructions.

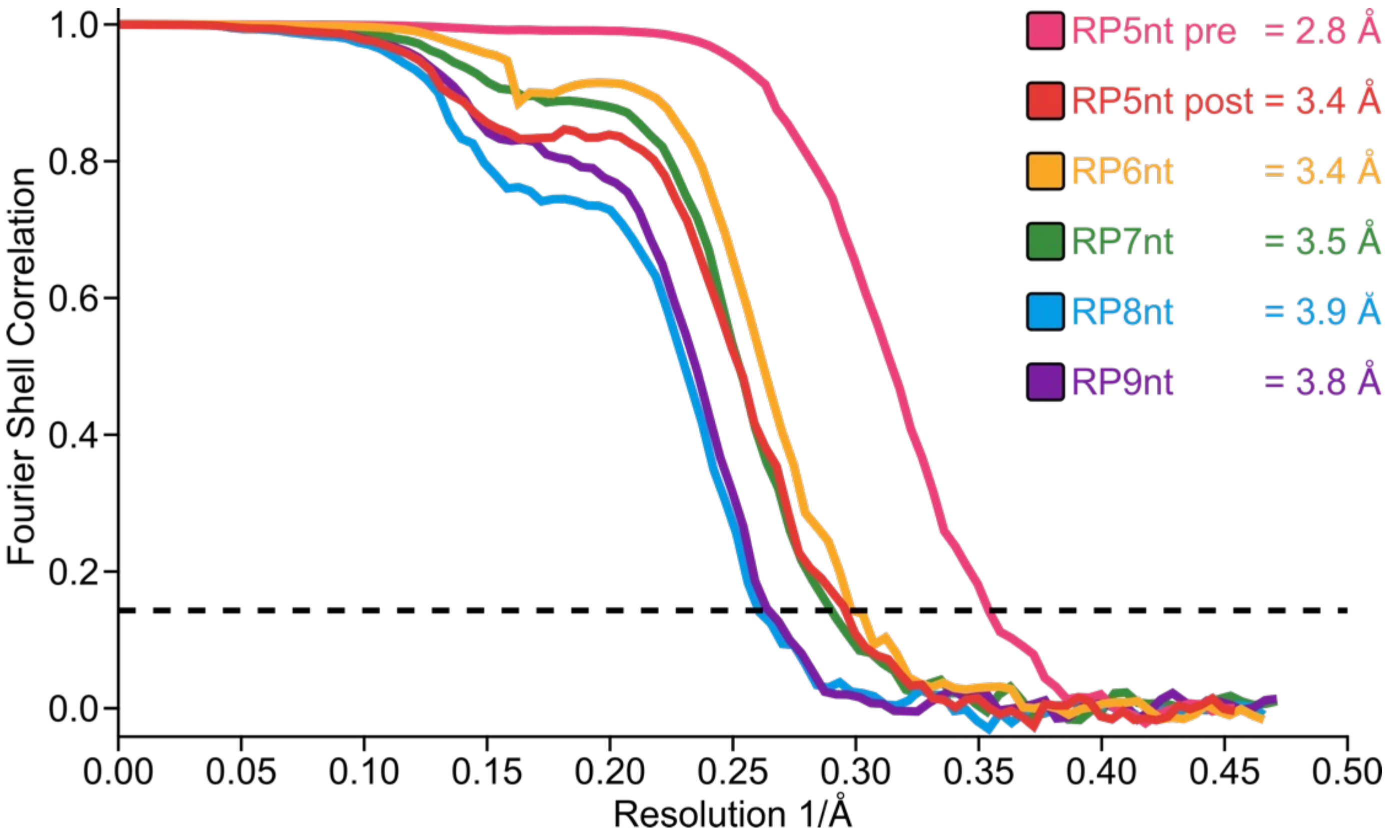


Supplementary Figure 3: Fourier shell correlation curves (based on corrected maps). Resolution indicated are based on 0.143 criterium

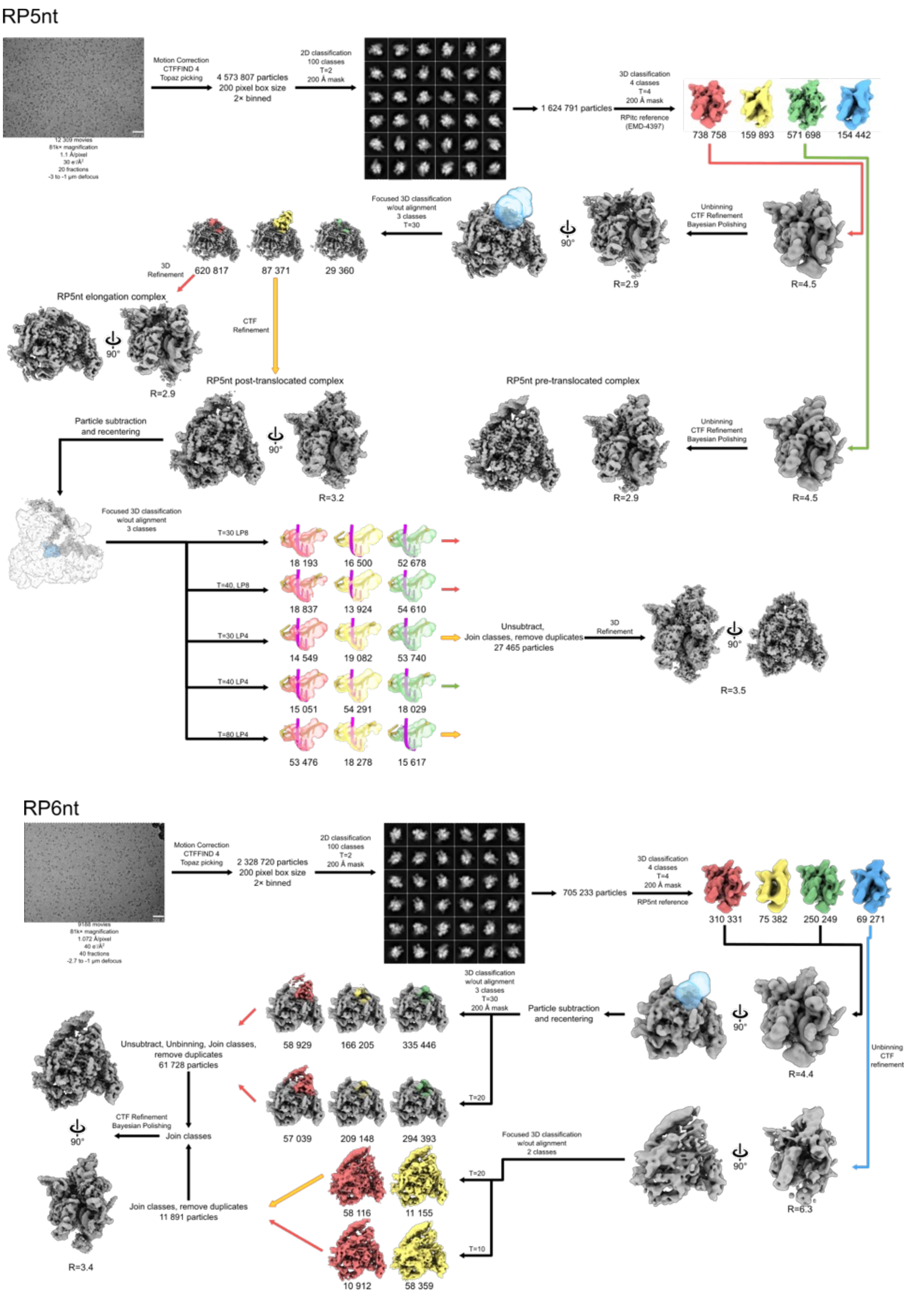


Supplementary Figure 4: Data processing pipeline for the RPitc 5nt dataset. Colour of arrows indicates the classes chosen.

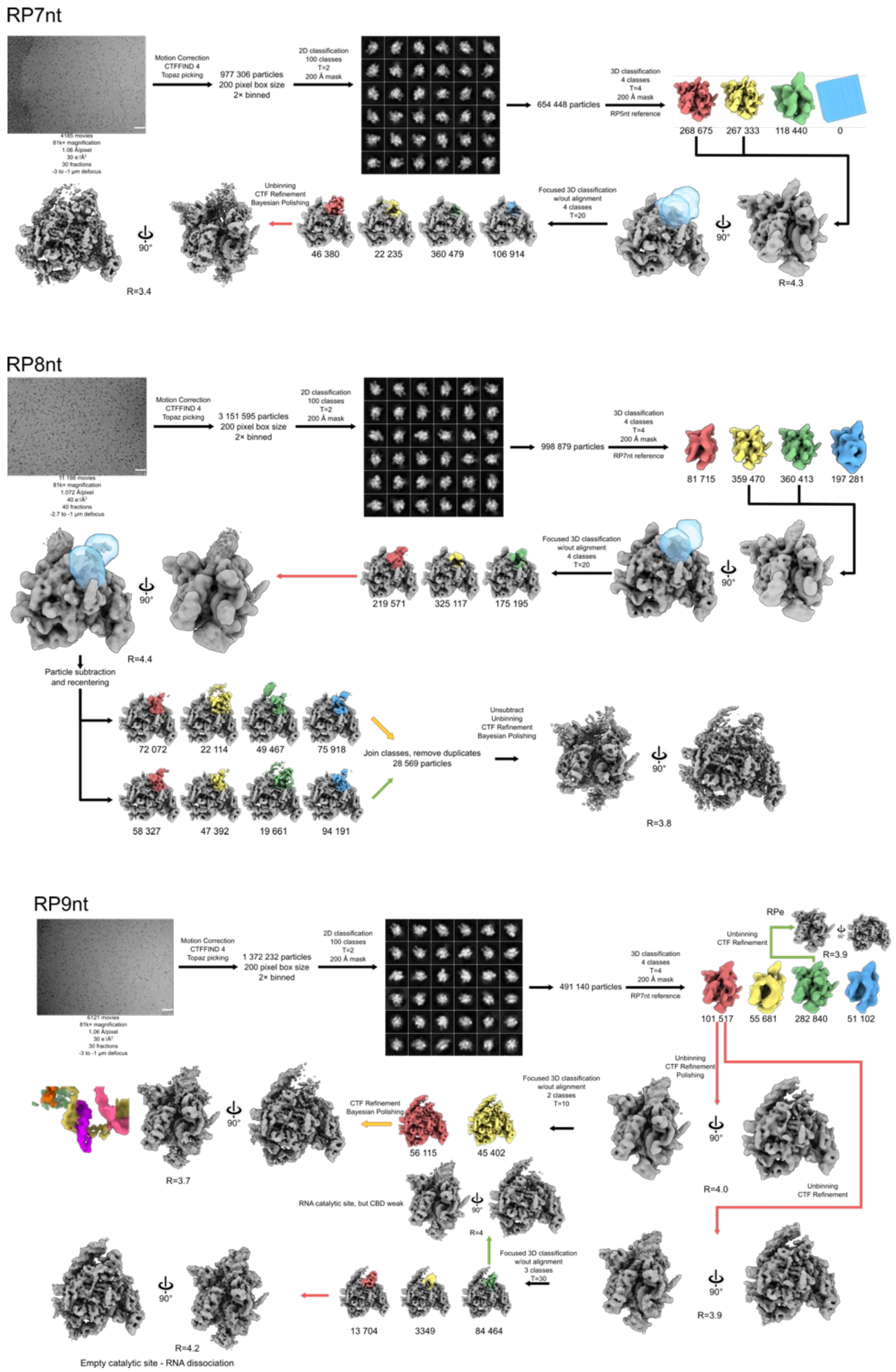


Supplementary Figure 5: Data processing flowcharts for 7-9nt RNA. Colour of arrows indicates the classes chosen.

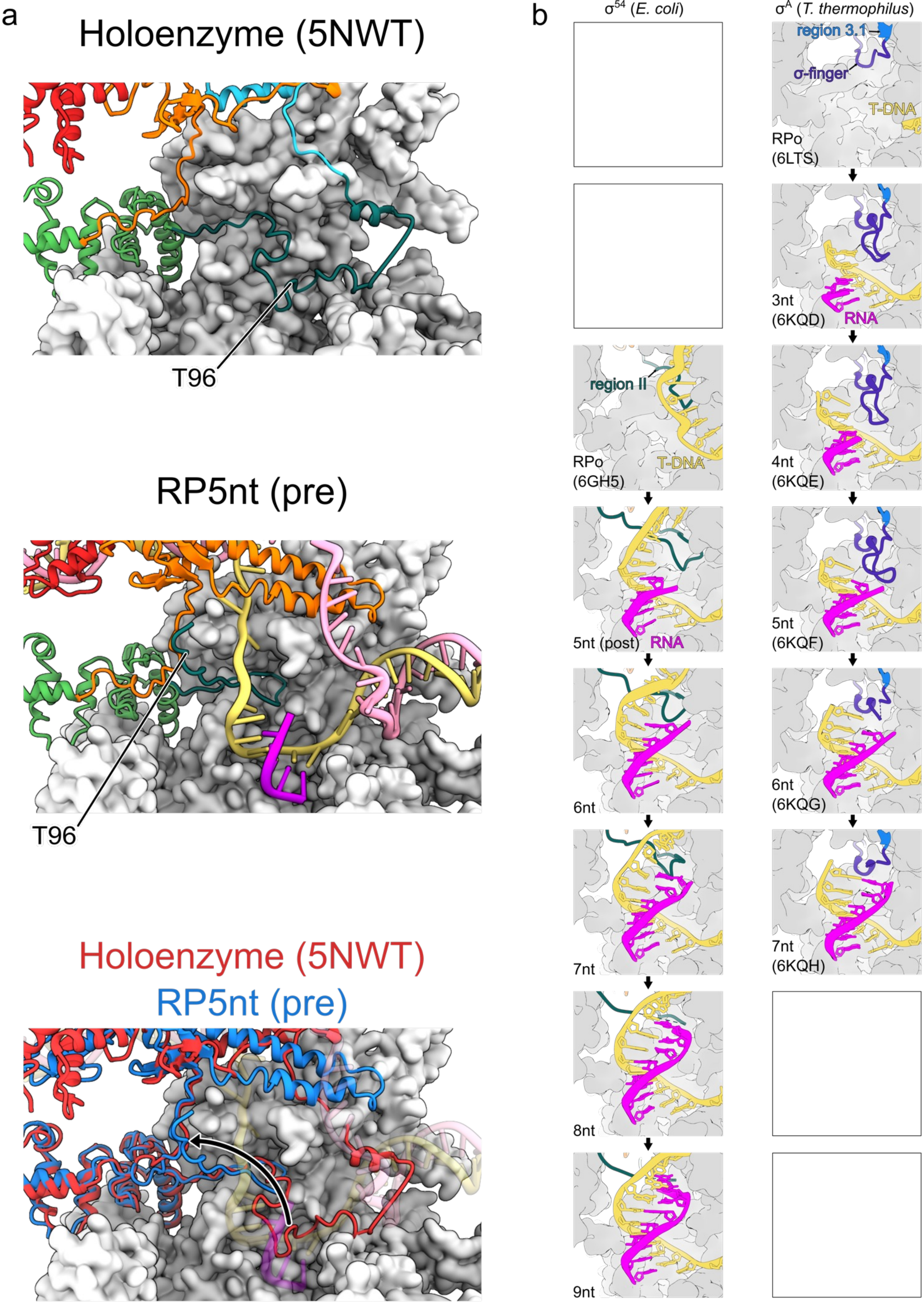


Supplementary Figure 6: RII finger conformations. (b) Comparison of RII in holoenzyme (5NWT) and RP5nt (pre) structure showing the relocation of RII upon binding the DNA-RNA hybrid. (b) Comparison of $\sigma^{54}$ and $\sigma^{70}$ ($\sigma^{A}$) initially transcribing complexes demonstrates that both utilise a similar mechanism of displacement of subdomains blocking the RNA exit channel, in which the 5' end of the growing RNA displaces σ factor, causing it to fold backwards..
